# Epigenomic reprogramming of repetitive noncoding RNAs and IFN-stimulated genes by mutant KRAS

**DOI:** 10.1101/2020.11.04.367771

**Authors:** Roman E. Reggiardo, Sreelakshmi Velandi Maroli, Haley Halasz, Mehmet Ozen, David Carrillo, Erin LaMontagne, Lila Whitehead, Eejung Kim, Shivani Malik, Jason Fernandes, Georgi Marinov, Eric Collisson, Utkan Demirci, Daniel H. Kim

## Abstract

RAS genes are the most frequently mutated oncogenes in cancer. However, the effects of oncogenic RAS signaling on the noncoding transcriptome are unclear. We analyzed the transcriptomes of human airway epithelial cells transformed with mutant KRAS to define the landscape of KRAS-regulated noncoding RNAs. We found that oncogenic KRAS upregulates noncoding transcripts throughout the genome, many of which arise from transposable elements. These repetitive noncoding RNAs exhibit differential RNA editing in single cells, are released in extracellular vesicles, and are known targets of KRAB zinc-finger proteins, which are broadly down-regulated in mutant KRAS cells and lung adenocarcinomas. Moreover, mutant KRAS induces IFN-stimulated genes through both epigenetic and RNA-based mechanisms. Our results reveal that mutant KRAS remodels the noncoding transcriptome through epigenomic reprogramming, expanding the scope of genomic elements regulated by this fundamental signaling pathway and revealing how mutant KRAS induces an intrinsic IFN-stimulated gene signature often seen in ADAR-dependent cancers.

## INTRODUCTION

Most of the human genome is noncoding and transcribed into RNA(*1*). About half of the human genome is comprised of transposable elements (TE) (*2*), whose expression patterns are often altered in cancer (*3*). Additionally, TEs contribute substantially to the noncoding transcriptome and are present in the exonic sequences of thousands of long noncoding RNAs (lncRNAs) (*4, 5*). Noncoding RNAs become disrupted in cancer (*6*) and epigenetic reprogramming (*7*), where activation of RAS signaling leads to repression of specific microRNAs (*8*) and coordinate regulation of lncRNAs in single cells, respectively. In lung cancers, RAS mutations are present in a third of lung adenocarcinomas (*9*) and serve as driver mutations that initiate tumorigenesis (*10*). Although RAS genes are among the most frequently mutated oncogenes in cancer (*11*), including pancreatic (*12*) and colorectal cancers (*13*), the extent to which oncogenic RAS signaling regulates the noncoding transcriptome remains unknown.

To investigate how mutant KRAS regulates the noncoding transcriptome during the initial stages of cellular transformation, we characterized the composition of protein-coding RNA, lncRNA, transposable-element derived repetitive noncoding RNA, and extracellular RNA, as well as global chromatin accessibility, using human airway epithelial cells (*14*) that have a constitutively active mutant KRAS(G12D) allele. We show that oncogenic KRAS induces cell-intrinsic interferon (IFN)-stimulated gene (ISG) signatures through epigenetic and RNA-mediated mechanisms, and that KRAB zinc finger (KZNF) genes that repress repetitive noncoding RNA loci are globally down-regulated both *in vitro* and *in vivo* in lung adenocarcinoma patients with activating mutations in KRAS. Our data reveal that significant upregulation and extracellular release of repetitive noncoding RNAs and ISGs are early transcriptomic signatures of mutant KRAS signaling.

## RESULTS

### Transcriptomic signatures of mutant KRAS

To determine the transcriptomic landscape of protein-coding and noncoding RNAs regulated by oncogenic RAS signaling, we performed RNA sequencing (RNA-seq) on human airway epithelial cells (AALE) that undergo malignant transformation upon introduction of mutant KRAS (Supplementary Figure 1a) (*14*). We compared the transcriptomes of AALE cells transduced with control lentiviral vector to AALEs that were transduced and transformed by mutant KRAS-containing lentiviral vector and performed differential expression analysis. We identified many protein-coding RNAs that were significantly differentially expressed (n=4323 upregulated, n=4711 down-regulated), as well as hundreds of differentially expressed lncRNAs (n=279 upregulated, n=409 down-regulated), revealing the broad extent to which mutant KRAS reprograms the transcriptome (Supplementary Figure 1b). For differentially expressed genes in mutant KRAS versus control AALEs, a large proportion of upregulated lncRNAs (∼75%) contained TE sequences in their exons, suggesting that repetitive noncoding loci in the genome are preferentially misregulated during mutant KRAS-mediated malignant transformation.

### Mutant KRAS induces an intrinsic IFN-stimulated gene signature

To explore the biological pathways that are perturbed by oncogenic RAS signaling, we performed gene set enrichment analysis (GSEA) (*15*) using genes that were differentially expressed in our mutant KRAS AALE cells. GSEA revealed that the most significantly enriched pathway was the interferon (IFN) alpha response, while the third most enriched pathway was the IFN gamma response (Supplementary Figure 1c). We examined all known ISGs that were expressed in our mutant KRAS AALEs and found that most of these genes were significantly upregulated, comprising a 25-gene mutant KRAS ISG signature (Fig. 1a). We then compared our KRAS ISG signature to IFN alpha (INFa) and IFN gamma (IFNg) response genes, along with the ADAR-dependent ISG signature described in Liu et al. (*16*). While the majority of KRAS ISG signature genes overlapped with the other 3 IFN-related signatures, several genes were unique to mutant KRAS cells (Fig. 1b). These results reveal that mutant KRAS signaling activates an intrinsic ISG response in transformed AALEs.

**Fig. 1.**
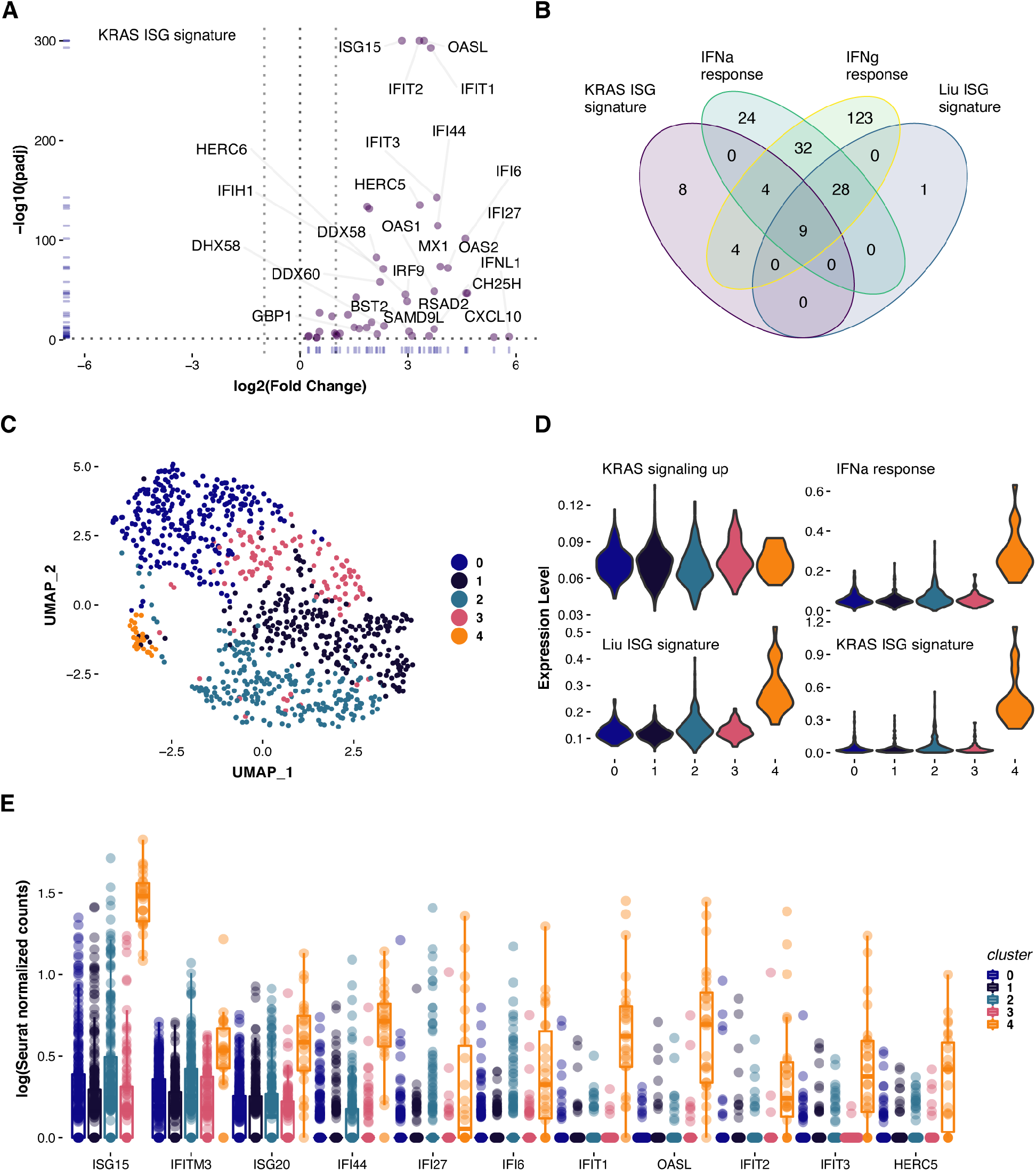
Mutant KRAS induces an intrinsic IFN-stimulated gene signature. **a**. Volcano plot depicting the differential expression of IFN-stimulated genes in mutant KRAS AALEs. **b**. Venn diagram highlighting the unique and shared genes across GSEA-enriched IFN gene sets. **c**. Seurat UMAP with k-means clusters representation of single cell RNA sequencing results. **d**. Violin plots depicting the distribution of averaged gene set expression in cells belonging to each k-means cluster. **e**. Distributions of normalized counts for DE ISGs across Seurat-identified clusters of single cells.

We then investigated whether this mutant KRAS ISG signature was specific to lung cells or if other cell types responded similarly. We performed RNA-seq on human embryonic kidney cells (HA1E) that were primed for oncogenic KRAS-driven transformation (*17*) and analyzed how mutant KRAS altered their transcriptomes. Similar to transformed AALEs, we also observed that thousands of protein-coding RNAs (n=2635 up, n=2639 down) and hundreds of lncRNAs were upregulated (n=165) or down-regulated (n=223) (Supplementary Figure 1d). When we performed GSEA, however, there was no enrichment for any IFN pathways in mutant KRAS-transformed HA1E cells, even though they were significantly enriched for upregulated KRAS signaling. In contrast to the mutant KRAS AALEs, we found that both IFNg and IFNa response pathways were among the most significantly decreased gene sets in mutant KRAS HA1Es (Supplementary Figure 1e), highlighting the cell type-specific differences in how the transcriptome is reprogrammed by mutant KRAS.

To assess the heterogeneity of the KRAS ISG signature in mutant KRAS AALEs, we performed single-cell RNA-seq (scRNA-seq) (n=1503 cells) (Fig. 1c). While KRAS signaling genes were upregulated in each cluster of cells, there was extensive heterogeneity in the IFNa response, Liu ISG, and KRAS ISG signatures, with the highest levels present in cluster 4 (Fig. 1d). This was also consistent at the gene level for ISGs (Fig. 1e), and these results indicate that individual cells respond differently to oncogenic KRAS signaling.

### Epigenetic reprogramming of ISGs by mutant KRAS

To elucidate the molecular mechanisms involved in inducing intrinsic ISG signatures in mutant KRAS AALE cells, we performed Assay for Transposase-Accessible Chromatin using sequencing (ATAC-seq) (*18*). In mutant KRAS AALEs, open chromatin was strongly enriched at gene promoters for upregulated KRAS signaling genes, as well as KRAS ISG signature genes (Fig. 2a). Open chromatin peaks were enriched at transcriptional start sites (TSS) of the ISGs OAS1, CH25H, GBP1, and IFNL1, all of which were significantly and specifically upregulated by mutant KRAS signaling (Fig. 2b). We found that several of these KRAS ISGs, including OAS2 and IFIT1, contained interferon-stimulated response elements (ISRE) that are recognized by the transcription factor IRF9, which also undergoes epigenetic activation in mutant KRAS AALEs (Fig. 2c). Some of the most enriched motifs at ATAC-seq peaks in mutant KRAS cells were for IRF9, IRF1, and AP-1 (FOS) (Fig. 2d), and genes at these open loci were significantly upregulated, as were the transcription factors that bind to these enriched motifs (Fig. 2e). These results reveal that oncogenic KRAS signaling induces intrinsic ISG expression through the epigenetic activation of ISG promoters.

**Fig. 2.**
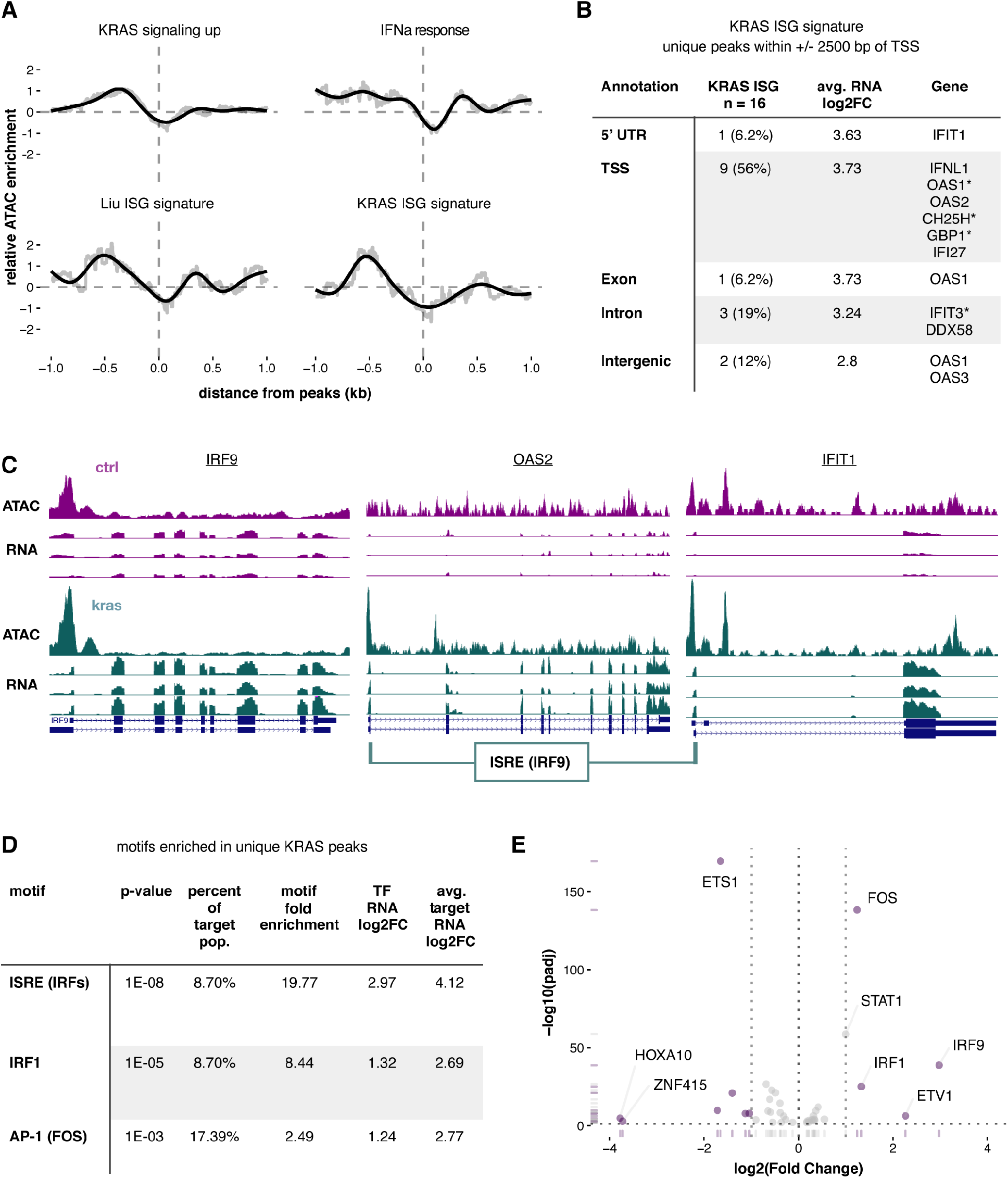
Epigenetic reprogramming of ISGs by mutant KRAS. **a**. Relative ATAC-seq peak density (KRAS/CTRL) calculated proximal to the TSS of genes within each respective gene set. **b**. Annotations of ISG ATAC-seq peaks unique to KRAS AALEs. * indicates 2 unique peaks. **c**. UCSC genome browser tracks of ATAC-seq and RNA-seq alignments in both KRAS and CTRL AALEs for IRF9, OAS2, and IFIT1. **d**. Selected enriched motifs from the unique KRAS ATAC-seq peaks assigned to ISGs by Homer. **e**. Volcano plot of all differentially expressed transcription factors with motifs in unique KRAS ATAC peaks assigned to ISGs by Homer.

### Mutant KRAS upregulates transposable element-derived noncoding RNAsh

We next investigated the molecular basis for intrinsic ISG signature activation in mutant KRAS AALE cells by analyzing the abundance of repetitive noncoding RNAs transcribed from TEs, which induce an IFN response in cancer cells when aberrantly expressed (*19, 20*). The LINE-1 element L1MC4a, the Alu elements AluSx, AluSg, AluJo, AluY, and AluSz6, and the hAT-Charlie element MER20 were all significantly upregulated in mutant KRAS AALE cells (Fig. 3a), suggesting that oncogenic KRAS signaling induces an ISG signature in transformed lung cells through the activation of TE-derived noncoding RNAs. We examined TE expression heterogeneity in our single-cell RNA-seq data from mutant KRAS AALEs and did not observe substantial heterogeneity in ALU, LINE, MER, or LTR class TE expression (Fig. 3b), as well as for the specific TEs L1MC4a and AluSx (Fig. 3c). When we performed RNA editing analysis on the scRNA-seq data to look for A-to-I editing (*21*), however, we found that TE-derived double-stranded RNAs (dsRNAs) exhibited significantly lower levels of RNA editing in cluster 4 (Fig. 3d), even though TE RNA levels were similar across all of the scRNA-seq data clusters. Mutant KRAS cells with lower levels of RNA editing exhibited higher expression of the dsRNA sensors MDA5 and RIG-I, while all of the clusters showed relatively similar levels of PKR expression, with a slightly higher level of PKR expression in the cells with lower RNA editing (Fig. 3e). These results reveal the extent of single cell heterogeneity in both RNA editing and dsRNA sensor ISGs in mutant KRAS cells.

**Fig. 3.**
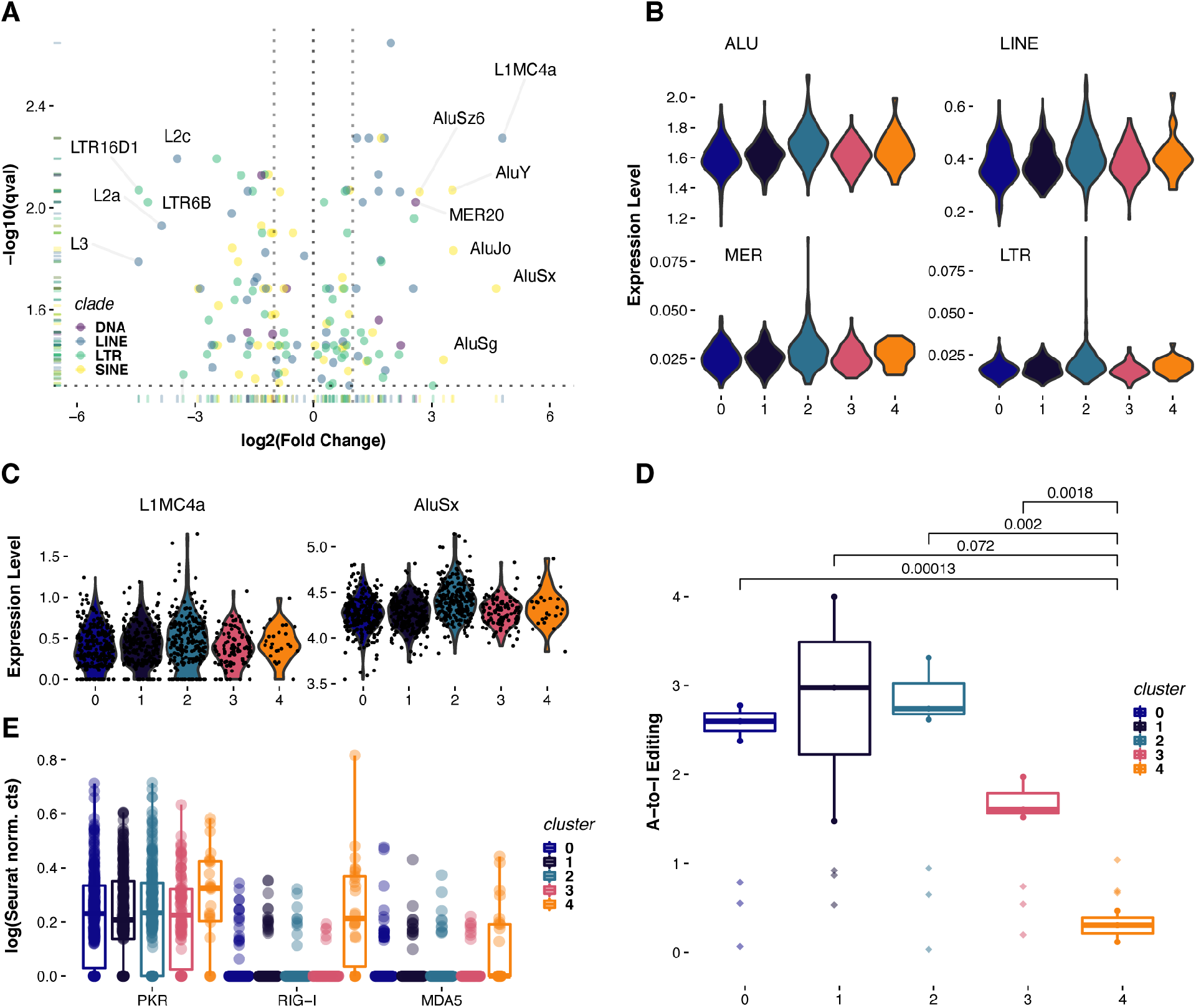
Mutant KRAS regulates repetitive noncoding RNA expression and editing. **a**. Volcano plot of all differentially expressed TE species aggregated from individual loci. **b**. Violin plots of the distribution of TE family expression across each single cell cluster. **c**. Violin plots depicting distribution of L1MC4a and AluSx expression across single cells clusters. **d**. Boxplots of Alu RNA editing from equal subsamples of cells from each single cell cluster. **e**. Distribution of normalized counts for PKR, MDA5, and RIG-I across Seurat-identified clusters.

To test whether extracellular RNAs that are released from mutant KRAS cells might also exhibit differential RNA editing, we isolated extracellular vesicles from the culture media of control and mutant KRAS AALEs (*22, 23*). Extracellular vesicles isolated from mutant KRAS AALE cell culture media were comprised of two different sized classes of vesicles that were ∼150nm and ∼213nm in diameter, while vesicles from control AALE media were ∼196nm in size (Supplementary Figure 2a). We performed RNA-seq and found that extracellular vesicles were enriched in lncRNAs when compared to the intracellular RNA composition (Supplementary Figure 2b). We also observed strong correlation between differentially expressed genes in intracellular and extracellular RNA-seq data (Supplementary Figure 2c). Moreover, repetitive noncoding RNAs derived from Alu, ERV, and L1 elements were significantly enriched in extracellular vesicles (Supplementary Figure 2d) but did not exhibit differential RNA editing (Supplementary Figure 2e), suggesting that intrinsic KRAS ISG signatures in mutant KRAS AALEs are not significantly affected by TE-derived repetitive noncoding RNAs that are packaged into extracellular vesicles.

### Broad epigenetic silencing of KRAB zinc-finger genes by mutant KRAS

Given the known roles of KRAB zinc-finger proteins (KZNFs) in TE silencing (*24*), we examined whether KZNFs were involved in TE regulation in mutant KRAS AALEs. When we examined the differential expression of KZNFs in mutant KRAS AALEs, we observed a broad and significant down-regulation of repressive KRAB domain-containing zinc-finger proteins (Fig. 4a) (Supplementary Figure 3a). Based on our ATAC-seq experiments, we determined that significantly down-regulated KZNF genes exhibited loss of open chromatin at their TSS and intronic regions (Fig. 4b), revealing that mutant KRAS signaling induces epigenetic silencing of many KZNF genes. Several significantly down-regulated KZNFs, including ZNF90, ZNF736, and ZNF683, were enriched for motifs in their TSS regions for ETS and ELK transcription factors (Fig. 4c), which are known downstream effectors of the RAS signaling pathway together with AP-1 (FOS) (*11*) (Fig. 4d).

**Fig. 4.**
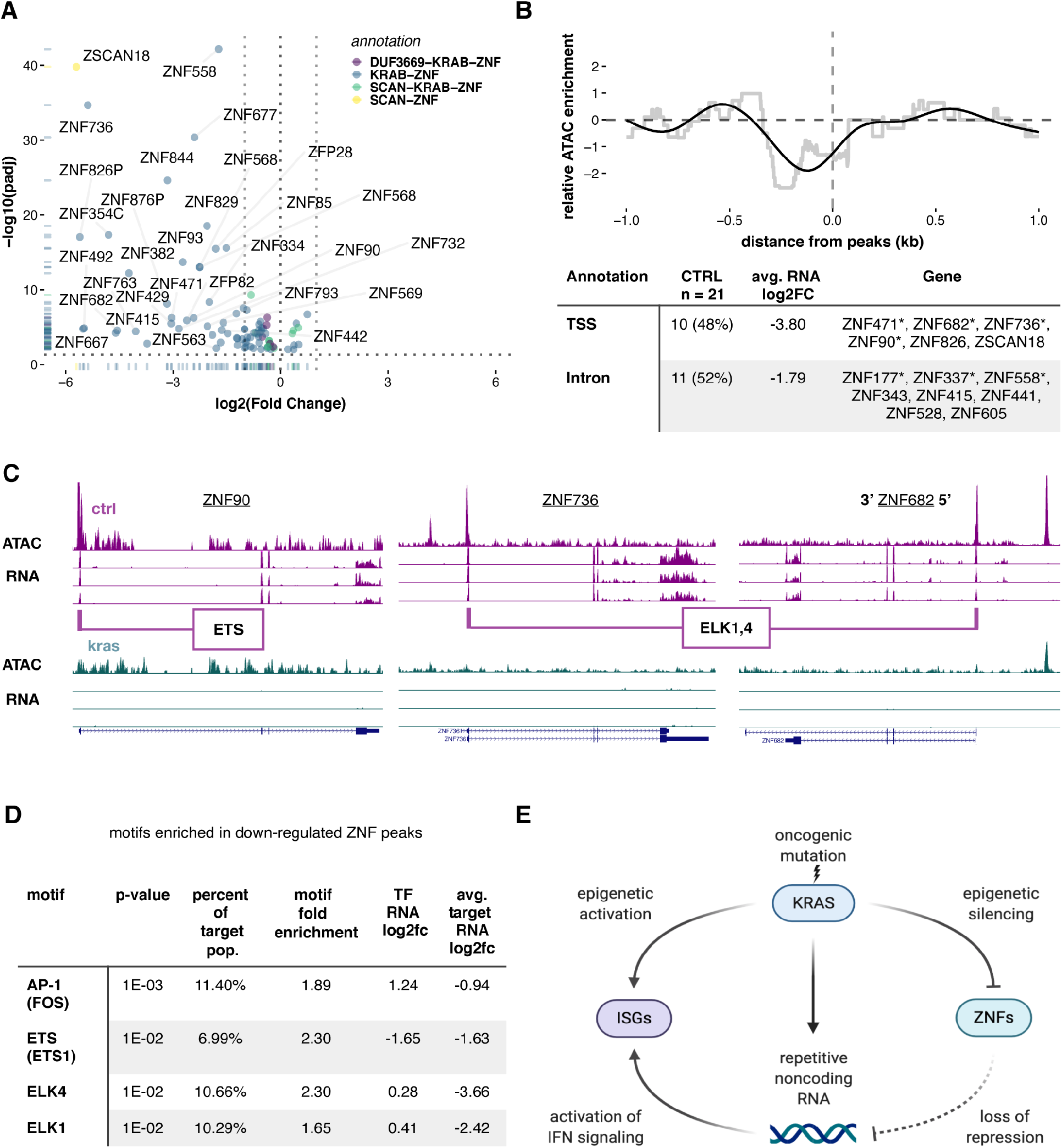
Broad epigenetic silencing of KRAB zinc-finger genes by mutant KRAS. **a**. Volcano plot of all differentially expressed zinc-finger genes with known annotations. **b**. Relative ATAC-seq peak density (KRAS/CTRL) calculated proximal to the TSS of KZNF genes and annotations of ATAC-seq peaks assigned to KZNFs and unique to CTRL AALEs. * indicates 2 unique peaks. **c**. UCSC genome browser tracks of ATAC-seq and RNA-seq alignments in both KRAS and CTRL AALEs of down-regulated ZNF90, ZNF736, and ZNF682. **d**. Selected enriched motifs from the unique KRAS ATAC-seq peaks assigned to KZNFs by Homer. **e**. Model of epigenomic reprogramming of KZNFs, repetitive noncoding RNAs, and IFN-stimulated genes by mutant KRAS (created with BioRender.com).

We then analyzed KZNF chromatin immunoprecipitation sequencing (ChIP-seq) data (*24*) using the University of California Santa Cruz (UCSC) Repeat Browser platform (*25*). We found that several of the significantly down-regulated KZNFs in mutant KRAS AALEs bind to the consensus TE sequences of MER20 and L1MC4a elements (Supplementary Figure 3b), which are specifically and significantly upregulated in mutant KRAS AALEs (Fig. 3a). This suggests that suppression of these KZNFs via oncogenic RAS signaling leads to de-repression of TE-derived noncoding RNAs during cellular transformation. This model is supported by broad and significant down-regulation of these same KNZFs *in vivo* in mutant KRAS-driven lung adenocarcinomas (Supplementary Figure 3c).

## DISCUSSION

Collectively, our findings reveal the genomic impact of oncogenic KRAS signaling on repetitive noncoding RNAs and ISGs. Our conclusions are based on deeply sequencing and analyzing the transcriptomes of mutant KRAS-transformed AALE cells at the population, single-cell, and extracellular levels, as well as the epigenomic level, building on our previous work identifying noncoding RNAs that are coordinately regulated with RAS signaling genes during epigenomic reprogramming (*7*). The molecular basis for the intrinsic ISG signature we observe in mutant KRAS AALE cells is different from TE-induced IFN responses in cancer cells treated with DNA methyltransferase inhibitors (*19, 20*), as we instead find a prominent role for broad KZNF supression during early stages of mutant KRAS-driven cellular transformation. Our studies also suggest that oncogenic RAS signaling contributes to the early induction of intrinsic ISG signatures that are observed across many cancers and cancer cells lines with ADAR dependencies (*16, 26*). In summary, our studies show that mutant KRAS both directly and indirectly activates repetitive noncoding RNAs through activation of RAS pathway transcription factors and repression of KZNFs that target these TE-containing loci in the human genome. Moreover, mutant KRAS reprograms the epigenome to both directly and indirectly activate intrinsic ISG signatures through opening chromatin at ISG promoters and activating repetitive noncoding RNAs that are recognized by dsRNA-binding RNA sensors such as PKR and MDA5 (Fig. 4e). Notably, the enrichment of both lncRNAs and TE-derived repetitive noncoding RNAs in extracellular vesicles released from mutant KRAS cells highlights their potential utility as RNA-based liquid biopsy biomarkers for diagnosing RAS-driven cancers.

## MATERIALS AND METHODS

### Cell Lines

The AALE stable cell lines pBABE-mCherry Puro (control) and pBABE-FLAG-KRAS(G12D) Zeo (mutant KRAS) were generated using retroviral transduction, followed by selection in puromycin of zeocin, respectively, 2 days post-infection. Both lines were cultured in SABM Basal Medium (Lonza SABM basal medium) with added supplements and growth factors (Lonza SAGM SingleQuot Kit Suppl. & Growth Factors). AALE cell lines were maintained using Lonza’s Reagent Pack subculture reagents. The HA1E cell lines were generated using lentiviral transduction (pLX317) to generate control and mutant HA1E pLX317-KRAS(G12V) stable cell lines using puromycin selection, and cells were cultured in MEM-alpha (Invitrogen) with 10% FBS (Sigma) and 1% penicillin/streptomycin (Gibco). All cell lines tested negative for mycoplasma.

### siRNA Knockdowns

AALEs were seeded at 1×10^6^ cells per well of a 6-well plate in complete growth medium, then reverse transfected with 30pmol siRNA using RNAiMAX lipofectamine according to manufacturer’s protocol. Cells were grown for 3 days in transfection medium under standard culture conditions and then harvested for RNA isolation and qPCR as previously described.

### Cell Viability Assay

2×10^4^ cells were subtracted from each siRNA transfection well at the time of transfection and seeded into individual wells of an ultra-low adhesion 96-well plate. The cells were grown in standard culture conditions for 4 days. They were then harvested, and ATP production was measured using the Cell TiterGLO Luminescent Cell Viability Assay (Promega) following the manufacturer’s protocol. Luminescence was measured on a Perkin Elmer VICTOR light 1420 Luminescence Counter.

### RNA Isolation & Purification

For AALE cell lines, bulk RNA was isolated from cells using Quick-RNA MiniPrep kit (Zymogen). All RNA was quantified via NanoDrop-8000 Spectrophotometer. For HA1E cell lines, bulk RNA was isolated using RNeasy Mini Kit (Qiagen) and quantified via Qubit RNA BR assay kit (Thermo).

### qPCR

cDNA was transcribed from 1ug RNA using iScript cDNA Synthesis Kit (Bio-Rad) according to manufacturer protocol. cDNA was diluted 1:6 and run with iTaq Universal SYBR Green Supermix (Bio-Rad) on ViiA 7 Real-Time PCR System according to manufacturer protocol. Cycle Threshold (CT) values were converted using Standard analysis. Values obtained for target genes were normalized to HPRT.

### RNA-seq

For AALE cell lines, 1ug of total RNA was used as input for the TruSeq Stranded mRNA Sample Prep Kit (Illumina) according to manufacturer protocol. Library quality was determined through the High Sensitivity DNA Kit on a Bioanalyzer 2100 (Agilent Technologies). Multiplexed libraries were sequenced as HiSeq400 100PE runs. For HA1E cell lines, 1ug of total RNA was used for mRNA enrichment with Dynabeads mRNA DIRECT kit (Thermo). First strand cDNA was generated with AffinityScript Multiple Temperature reverse transcriptase with oligo dT primers. Second strand cDNA was generated with mRNA Second Strand Sythesis Module (New England Biolab). DNA was cleaned up with Agencourt AMPure XP beads twice. Qubit dsDNA High Sensitivity Assay was used for concentration measurement. 1ng of dsDNA was further subjected to library preparation with Nextera XT DNA sample prep kit (Illumina) per manufacturer instructions. Library size distribution was confirmed with Bioanalyzer (Agilent). Multiplexed libraries were sequenced as NextSeq500 75PE runs.

### Single-cell RNA-seq

For single cell RNAseq, 1×10^6^ cells were harvested and re-suspended in 1mL 1X PBS/0.04% BSA (1000 cells/ul) according to the cell preparation guidelines in the 10X Genomics Chromium Single Cell 3’ Reagent Kit User Guide. GEMs were generated from an input of 3,500 cells. We used the 10X Genomics Chromium Single Cell 3’ Reagent Kits version 2 for both the GEM generation and subsequent library preparation and followed the manufacturer’s reagent kit protocol. Quantification of all RNAseq libraries was performed by QB3 at UC Berkeley. RNAseq libraries were sequenced as HiSeq4000 100PE runs.

### ATAC-seq

100,000 cells were collected and centrifuged at 500xg for 5 minutes at 4C. Pellets were washed with ice-cold PBS and centrifuged. Pellets were resuspended in ice-cold lysis buffer. Tagmentation reaction and purification were conducted according to manufacturer’s protocol (Active Motif). Libraries were sequenced on a NextSeq500 as 2×75 paired end reads.

### Extracellular RNA

The exoRNeasy serum/plasma maxi kit (Qiagen) was used to isolate extracellular vesicles, which were quantified using Nanoparticle Tracking Analysis (Malvern, UK). 30 ml of cell culture supernatant was filtered to remove particles larger than 0.8 um. The filtrate was precipitated with kit buffer and filtered through a column to collect extracellular vesicles. These vesicles were then lysed with QIAzol® lysis reagent. Total RNA was isolated using the indicated phase separation method and used to make libraries for RNA-seq, which were sequenced on a NextSeq500.

### Exosomal RNA

Exosomes were isolated using the Exosome Total Isolation Chip (ExoTIC) as previously described (*23*). The ExoTIC device was first flushed with 2 mL of 1X PBS buffer. Then, the EVs from culture media were isolated as follows: a five milliliter-volume of culture medium was drawn up in the same syringe and connected with the ExoTIC device. This syringe along with the ExoTIC device, was fixed onto a syringe pump. A pump flow rate of 5 mL/h was applied to filter the culture media, concentrating EVs in front of the nanoporous membrane. Free proteins, nucleic acids, etc., which are smaller than the membrane pore size (~50 nm) pass through the filter pores. The EV-containing retentate was then washed by running 5 mL of 1X PBS through the device using the same syringe. The ExoTIC device was then disconnected from the syringe, and the purified EV solution was collected via the device inlet using a 200μL pipet. RNA extracted from the purified EV sample using the miRNeasy Mini Kit (Qiagen) was used to make libraries for RNA-seq, which were sequenced on a NextSeq500.

### Statistical Analysis

All quantitative data for functional assays has been reported as means ± standard deviation. Statistical; significance for these was calculated using a t-test and p-values <0.05 were considered significant.

### Data and Code Availability

All code for figures, file parsing, and data processing is available via https://github.com/rreggiar/aale.kras. Sequencing data is accessible via GEO accession GSE120566. Additionally, UCSC Genome Browser tracks used in figures is accessible at https://genome.ucsc.edu/s/rreggiar/kras.and.ctrl.atac.

### RNA-seq Analysis

All *fastq* files were trimmed with *Trimmomatic 2 (0*.*38)* (*27*) using the Illumina NextSeq PE adapters. The resulting trimmed files were assessed with *FastQC* (*28*) and then passed through the following analytical pipeline: *Salmon (1*.*0):* pseudoalignment of RNA-seq reads performed with *Salmon (29)* using the following arguments:

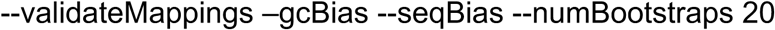

using an index created from the *GENCODE* version 32 transcriptome fasta file using standard arguments.

*STAR (2*.*7*.*3a):* trimmed reads were aligned to the Human genome with default arguments using a 2-pass approach described previously (*30*).

The resulting .sam files were converted to bam, sorted, and indexed using Samtools with default arguments for all procedures (*31*).

*Sleuth (0*.*30*.*0):* transcript differential expression was performed using *Sleuth (32)* and *Wasabi* (1.0.1) to convert the *Salmon* output into the proper format. Upon completion, the transcripts with q-values below 0.05 in the likelihood-ratio test were used to filter salmon output from which log2fc was manually calculated and paired to the sleuth output. Sleuth was primarily used for quantifying DE of Transposable Element loci in which case the provided reference was the repeat masked loci sequences from the UCSC Genome Browser.

*DESeq2 (1*.*24*.*0): Salmon* output was imported into a DESeq object using *tximport* (*33*) and differential expression analysis was performed with standard arguments (*34*). All results were filtered to have padj <= 0.01. In the case where R could only generate 0.00 for the padj values, they were reset to the lowest non-zero padj value in the data set. Where count data was used, it was normalized across samples using DESeq.

### Transposable Element Content Analysis

*Exon and 5’/3’ UTR Overlap:* a whole genome .*gtf* file was downloaded from the UCSC genome browser table browser utility. This file was parsed and merged with the GENCODE v.29 reference transcriptome. This modified .*gtf* (now a .*bed* file) was passed to *bedtools* (*35*) where the overlap function was used with the following arguments:

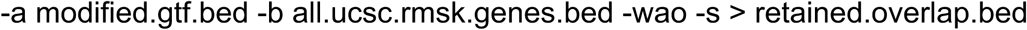

alongside a whole genome .*gtf* retrieved as described above except generated from the repeat-masked browser track. The resulting overlapped bed file was processed and visualized using custom *R* scripts.

#### Differential Expression

Differential transcript abundance was determined using the *Salmon* and *Sleuth* procedures described above provided with a custom index comprising both the *GENCODE* version 29 transcripts and all transcripts extracted from the Hammel lab GTF file as described in the single cell procedures. *Sleuth* output was filtered and visualized using *R* and *Tidyverse*.

### Zinc Finger Protein Analysis

ChIP-exo data and supplementary information were extracted from supplementary data provided by *Imbeault et al (24)*. ZNF genes were cross referenced with *DESeq2* and *RepeatMasker* (*36*) outputs to extract relevant differential expression data of ZNF proteins and Transposable Element transcripts using *R. RepeatMasker* output from promoter analyses was cross referenced with ChIP-exo target data to identify potential regulatory targets of differentially expressed KZNFs. Only KZNF targets with ‘score’ [see Imbeault *et al*] >= 75 were kept for analysis. Analysis of all data was performed and visualized in *R* using custom scripts.

### Gene Set Enrichment Analysis

Genes determined to be significantly differentially expressed in *DESeq2* output were first ‘pre-ranked’ in *R* by the following metric: Score metric = sin(log2FoldChange) * -log^10^(p-value)

The resulting ranked files objects were processed using the *R* package *fgsea* alongside gene set files downloaded from msigdb using the *R* package *msigdbr*. Additional code was written for select vizualizations.

### Gene Ontology Analysis

Upregulated gene names were extracted from *DESeq2* output using bash command line tools. Name lists were pasted into the *Gene Ontology Consortium*’s *Enrichment Analysis* tool powered by *PANTHER*. Output data was exported as .*txt* files and parsed using bash command line tools. Parsed data was visualized using custom *R* scripts.

### Single Cell Analysis

#### Cell Ranger

Single cell output data was processed using 10x pipeline CellRanger. The mkfastq functionality was used to generate fastq files for further downstream analysis.

#### Salmon – Alevin

fastq files generated above were passed to Salmon alevin with the default arguments for CHROMIUM V2 data. alevin was used to psuedoalign the libraries to both the GENCODE v32 combined with the repeat masked loci sequences extracted from Hg38 via the UCSC Genome Browser. A salmon index was built from this reference with standard arguments. These alevin output matrices were imported into R using tximport.

#### STAR-solo

This feature within the STAR software was used to generate single cell SAM files for downstream processing. Run with the recommended arguments. *Seurat (3*.*0):* Normalization and UMAP clustering were performed with Seurat following their described approach optimized to our data set (see code notebook). Additional code was written to extract count data from Seurat single cell objects using the SingleCellExperiment R package *(37)*.

### TCGA ZNF Analysis

TCGA-LUAD and GTEX lung phenotype and normalized count data were downloaded from the UCSC Xena browser TOIL data repository (https://xenabrowser.net/datapages/?cohort=TCGA%20TARGET%20GTEx&addHub=https%3A%2F%2Fxena.treehouse.gi.ucsc.edu&removeHub=https%3A%2F%2Fxena.treehouse.gi.ucsc.edu%3A443). The files were combined and patients were grouped by their KRAS mutation status and identity. These data were compared to and visualized alongside of data generated from our analysis using custom *R* code. Significance was determined with a one-way t test implemented in the *R t*.*test()* function.

### RNA Editing

#### RNAEditingIndexer

Single cell SAM files were subsampled into 3 equal subsets per cluster based on barcode. Each SAM file was then converted into a BAM as described above and used as input for the RNAEditingIndexer script with bed files generated by extracting the locations of detected Transposable Element loci from *Sleuth* output (*21*).

### ATAC-seq Analysis

#### ENCODE

The ENCODE ATAC-seq pipeline was used for alignment, quality control, MACS analysis with default arguments to produce output for downstream analysis.

#### HOMER

Narrow peaks files produced above were processed using a variety of HOMER tools with default arguments where not explicitly stated: *findMotifsGenome* was used to find enriched motifs in each ATAC library and their subsets.

*annotatePeaks* was used to generate detailed annotations of the motifs found above and their context. It was also used to create gene set differential accessibility histograms by using the -hist argument set to 1.

*mergePeaks* allowed us to identify unique and overlapping peaks across the two libraries. *makeTagDirectory* was used to generate peak height estimates to be applied on all histogram comparisons (*38*).

### Quantification and statistical analysis

All statistical analyses were performed with R (version 3.6.1) running from the Rocker ‘Tidyverse’ Docker container (rocker/tidyverse:3.6.1). Unpaired, bi-directional t test was performed with the t.test() function on samples with 3 biological or technical replicates. Linear regression was carried out with the lm() function.

### Additional Code

All analysis was performed in the *R* programming language with supplemental scripts written in *Bash*.

## ACKNOWLEDGEMENTS

We thank members of the Kim Lab, Brooks Lab, Haussler Lab, and Carpenter Lab for helpful discussions. This work was supported by funds from the Baskin School of Engineering and the Ken and Glory Levy Fund for RNA Biology (to D.H.K.), the NHGRI-funded UCSC Genomic Sciences Graduate Training Program (NIH T32 HG008345) (to R.E.R and H.H.), the NIGMS-funded UCSC IMSD Program (NIH R25 GM058903) (to D.C.), and the NCI (NIH R01 CA227807, NIH R01 CA239604, NIH R01 CA230263) (to E.C.).

## AUTHOR CONTRIBUTIONS

D.H.K. conceived and designed the study and wrote the manuscript, R.E.R. performed computational analysis and generated figures, S.V.M., H.H., M.O., D.C., E.L., L.W., E.K., and S.M. performed experiments, J.F., E.C., and U.D. provided resources, and G.M. performed initial computational analysis. The authors declare no competing interests.

## SUPPLEMENTARY FIGURES

**Supplementary Figure 1.**
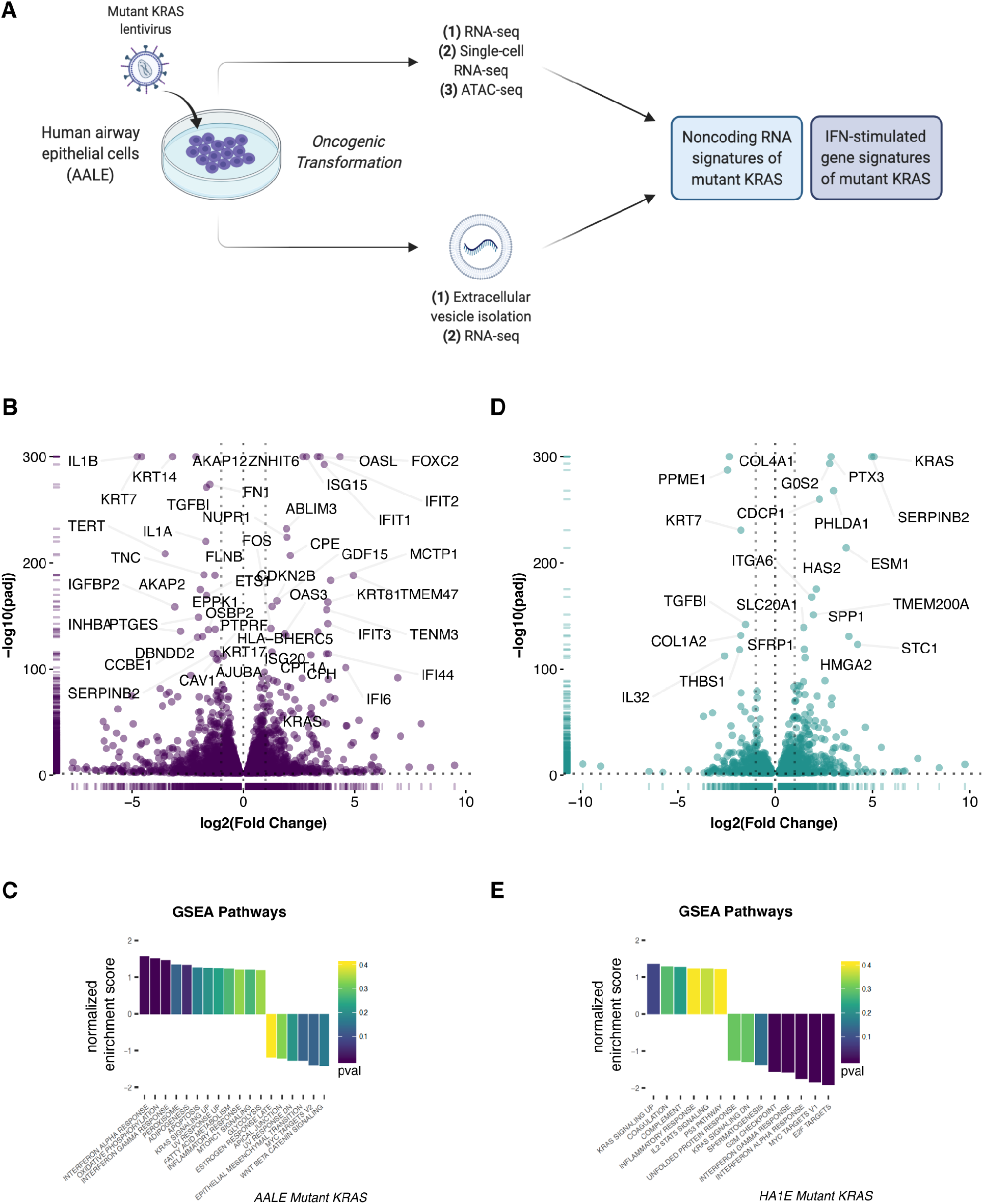
Mutant KRAS reprograms the transcriptome. **a**. Schematic of experimental workflow (created with BioRender.com). **b**. Volcano plot depicting the differential expression of GENCODE genes in mutant KRAS AALEs. **c**. GSEA normalized enrichment score for differentially expressed genes in mutant KRAS AALEs. **d**. Volcano plot depicting the differential expression of GENCODE genes in mutant KRAS HA1Es. **e**. GSEA normalized enrichment score for differentially expressed genes in mutant KRAS HA1Es.

**Supplementary Figure 2.**
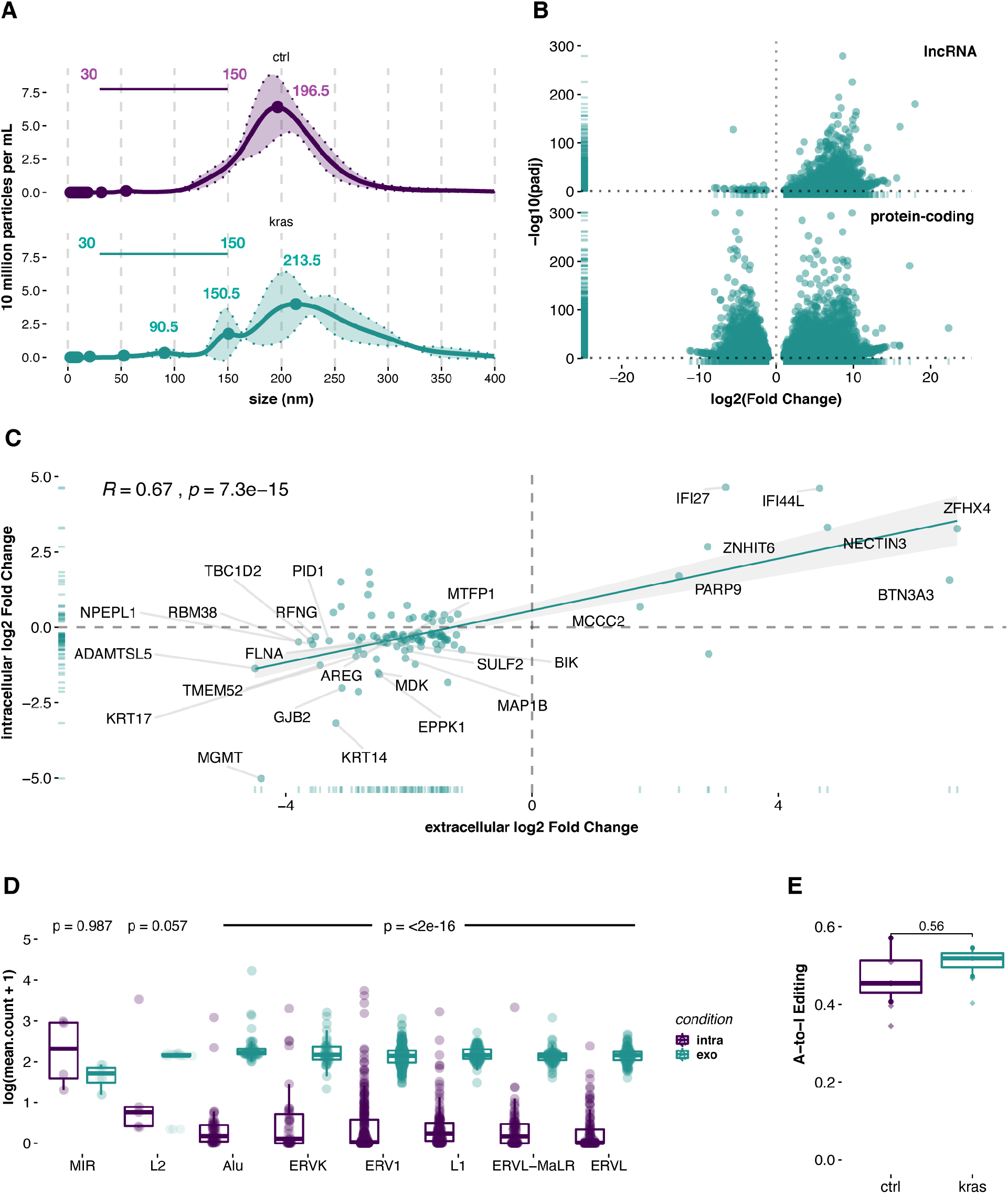
Extracellular RNAs released from mutant KRAS AALEs. **a**. Ribbon plot of extracellular vesicle size abundance from nanoparticle tracking analysis. Solid line is mean of three measurements and the surrounding filled-in area reflects their standard deviation. Solid points are local maxima with corresponding value labelled adjacently. **b**. Volcano plots depicting the differential enrichment of lncRNA and protein-coding genes in mutant KRAS AALE extracellular RNAs. **c**. Correlation between intracellular and extracellular RNAs that are differentially expressed/enriched from mutant KRAS AALEs. **d**. Boxplot comparing distributions of average counts across conditions in TE clades, significance determined by one-way t-test. **e**. Boxplot of detected Alu editing in three replicates of RNA-seq performed on extracellular vesicles isolated from media from each condition.

**Supplementary Figure 3.**
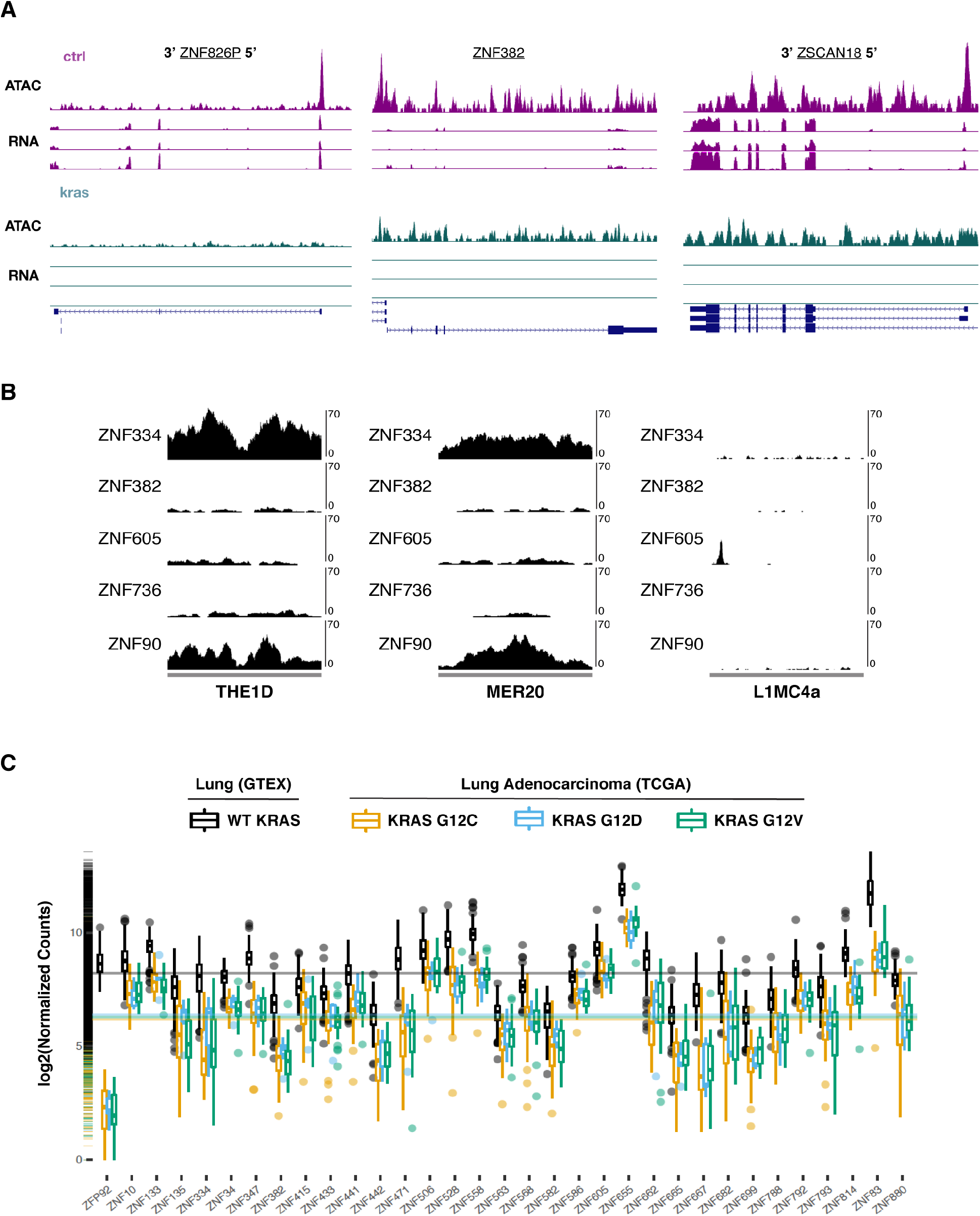
Mutant KRAS represses KZNFs *in vitro* and *in vivo*. **a**. UCSC genome browser tracks of ATAC-seq and RNA-seq alignments in both KRAS and CTRL AALEs of down-regulated ZNF826P, ZNF382, and ZSCAN18. **b**. UCSC repeat browser tracks of ChIP-seq peaks for differentially expressed KZNFs over indicated TE consensus sequences. **c**. Distribution of differentially expressed ZNFs (in mutant KRAS AALEs) in RNA-seq data from GTEx lung tissues and TCGA lung adenocarcinomas by KRAS mutation status.

